# BOA: A Partitioned View of Genome Assembly

**DOI:** 10.1101/2022.05.22.492973

**Authors:** Priyanka Ghosh, Xiaojing An, Patrick Keppler, Sureyya Emre Kurt, Ümit V. Çatalyürek, Sriram Krishnamoorthy, P. Sadayappan, Aravind Sukumaran Rajam, Ananth Kalyanaraman

## Abstract

*De novo* genome assembly is a fundamental problem in computational molecular biology that aims to reconstruct an unknown genome sequence from a set of short DNA sequences (or *reads*) obtained from the genome. High throughput sequencers could generate several billions of such short reads in a single run. However, the relative ordering of the reads along the target genome is *not* known *a priori*. This lack of information is one of the main contributors to the increased complexity of the assembly process. Typically, state-of-the-art approaches produce an ordering of the reads toward the *end* of the assembly process, making it rather too late to benefit from the ordering information. In this paper, with the dual objective of improving assembly quality as well as exposing a high degree of parallelism for assemblers, we present a partitioning-based approach. Our framework—which we call `BOA` (for bucket-order-assemble)—uses a bucketing alongside graph- and hypergraph-based partitioning techniques to produce a partial ordering of the reads. This partial ordering enables us to divide the read set into disjoint blocks that can be independently assembled in parallel using any state-of-the-art serial assembler of choice. We tested the BOA framework on a variety of genomes. Experimental results show that the hypergraph variant of our approach, `Hyper-BOA`, consistently improves both the overall assembly quality and performance. For the inputs tested, the `Hyper-BOA` framework consistently improves the N50 values of the popular standalone `MEGAHIT` assembler by an average of 1.70× and up to 2.13×; while the largest alignment length improves 1.47× on average and up to 1.94×. The time to solution also consistently improves between 3-4× for the system sizes tested.

## 1 Introduction

In *de novo* genome assembly, the relative ordering and orientation of the input reads along the target genome is not known *a priori*. In fact it can be argued that one of the primary contributors to the problem complexity is the lack of this information—i.e., if the ordering and orientation of the reads is known at input then the genome assembly problem would reduce to a simpler (albeit less exciting) problem of performing a linear sequence of pairwise alignments between adjacent reads to produce the assembly. However, the DNA sequencers preserve neither the genomic coordinates from where the reads were sequenced nor any significant relative ordering information between the reads (except for paired end read information). Consequently, assembly algorithms are left to infer an ordering and orientation along the course of their respective computations.

Different assembly approaches vary on how much they rely on the read ordering and orientation (henceforth abbreviated as *OO* for simplicity) information, and at what stages of their algorithm they try to infer it. De Bruijn graph assemblers [4, 17, 21], which now represent a dominant segment of modern day short-read assemblers, use an approach that is largely oblivious to OO information. This is because these assemblers use de Bruijn graphs that break the reads into shorter fixed-length k-mers at the early stages of the algorithm. Therefore, the information on how the reads are ordered/oriented along the target genome is typically not recoverable until the end of the assembly pipeline (i.e., until after contigs are generated). On the other hand, the more traditional overlap-layout-consensus (OLC) class of assemblers [16, 17, 22] are more explicit in trying to infer the OO information in their assembly pipeline—as the overlap phase aligns reads against one another with an intent to arrive at a read layout. And yet, because the overlap phase is also the most time consuming step of the assembly pipeline for the OLC assemblers, the OO information is practically not available until later stages of the assembly.

In this paper, we ask the simple question of what if either a total (ideal but not practical) or at least a partial order information can be generated earlier in the assembly computation^4^. Could that help improve performance and/or assembly quality? If so, what are some of the ways to generate such OO information earlier in the assembly algorithmic stages and what are their assembly efficacies?

### Contributions

To address the above questions, we present a parallel assembly framework that uses a graph partitioning-centric approach. Graph partitioning [8] is a classical optimization problem in graph theory that aims to partition the set of vertices of an input graph into a pre-determined number of partitions in a load balanced manner. The problem has seen decades of research in development and application under numerous contexts including in the parallel processing of graph workloads [10], as well as partitioning assembly graphs [19] and read datasets [1, 12].

In this paper, we exploit graph partitioning and its properties to produce a partial ordering of reads and in the process also enable parallelization of the assembly workload. More specifically:

- We cast the assembly problem in two forms: a) one that uses graph partitioning, and b) another that uses hypergraph partitioning.
- To enable the application for different types of partitioning, we propose a light-weight bucketing algorithm that bins reads into buckets based on fixed-length exact matches and uses the bins to generate graph/hypergraph representations suitable for partitioning.
- Once bucketed and partitioned, each individual part can be independently assembled. This strategy allows the user to use any standalone (off-the-shelf) assembler of choice. Consequently, we call our assembly framework `BOA` (stands for bucket-order-assemble). Two implementations (i.e., concrete instantiations) of this framework are presented and evaluated—one that uses a classical graph partitioner (ParMETIS [13]) and another that uses a hypergraph partitioner (Zoltan [5]).
- To comparatively assess the assembly efficacy of the partitioning-based approach, we also construct a benchmark *Oracle* assembly workflow that uses the correct read ordering available from sequencing simulators.

Experimental results demonstrate that our partitioning-based implementations a) improve parallel performance of assembly workloads; and b) improve assembly quality, consistently under several qualitative measures. In fact the partitioning-based approaches yield results that come closest in terms of quality to the Oracle assemblies produced.

The rest of this paper is organized as follows. In Section 2, we present relevant preliminaries about graph partitioning and hypergraph partitioning. In Section 3, we present our `BOA` assembly framework including details of our graph/hypergraph-based formulations. Section 4 presents an experimental evaluation of the `BOA` framework, comparing it with standalone short read assemblers as well as the benchmark results. Section 5 concludes the paper.

## 2 Preliminaries and Notation

### 2.1 Strings and Genome Assembly

Let *s* denote an arbitrary string over a fixed alphabet Σ, and let |*s*| denote the length of the string. Let *s*[*i, j*] denote the substring of *s* starting at index *i* and ending at index *j*. As a convention, we index strings from 1, and the *i^th^* character of *s* is denoted by *s*[*i*]. A *k*-mer is a (sub)string of length *k*.

Given a substring *s*[*i, j*] of *s*, we refer to the character immediately preceding the substring in *s* to be its “left-character” or *lchar* (if one exists). More specifically, *lchar_i_* = *s*[*i* − 1] if 1 < *i* ≤ |*s*|, and if *i* = 1, then *lchar_i_* = *B*, where *B* ∉ Σ is used to represent a blank symbol.

The input to genome assembly is a set of *n* reads (denoted by 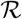). Each read is a string over the alphabet Σ = {*a, c, g, t*}. We denote the reverse complemented form of a read *r* as *rc*(*r*). If reads are generated with paired-end information, then the two reads of the same pair are assigned consecutive read IDs *i* and *i* + 1, so that the odd read ID corresponds to the forward strand read and the even read ID corresponds to the reverse strand read. We denote the set of all forward (alternatively, reverse) reads as 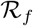 (alternatively, 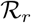). Note that 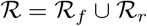, and 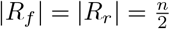.

### 2.2 Graph Partitioning

A undirected *graph* 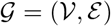 is defined by a set of vertices 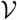 and a set of edges 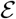. An edge *e_i,j_* is a pair of distinct vertices, i.e. *e_ij_* = {*v_i_, v_j_*}, 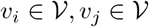. The degree *d_i_* of a vertex *v_i_* is defined as the number of edges incident to that vertex. Weights and costs can be assigned to vertices and edges. 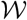 is used to represent the weight assignment for vertices, where *w_i_* is the weight for the vertex 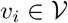. 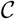 is the cost assignment for edges, where *c_ij_* represents the cost for the edge 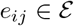.

A *K*-way partition of 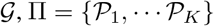, places each vertex of the graph into a *part*. More concretely, Π is a *K*-way partition if each part 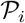 is a non-empty subset of 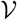, each pair of parts is disjoint, i.e., 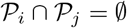 for all 1 ≤ *i* ≠ *j* ≤ *K*, and the union of all parts recovers 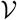, that is 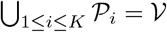. For a *K*-way partition Π, an edge *e_ij_* = { *v_i_, v_j_*} is called external (or *cut*) if 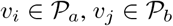 with *a* ≠ *b*, otherwise called internal (or *uncut*). 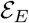 is used to represent the set of all external edges. The cost (or *cutsize*) *χ* of Π is defined as: 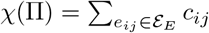. A *K*-way partition, Π, is called balanced if the following holds:

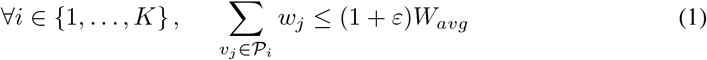

where, 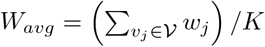, and *ε* is a given maximum imbalance ratio.

The graph partitioning problem is defined as follows: given a graph 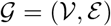, vertex weight and edge cost assignments 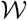 and 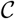, a part number requirement *K*, and the maximum allowed imbalance ratio *ε*, find a balanced *K*-way partitioning that minimizes the cost. The graph partitioning problem is known to be NP-hard [7], even for seemingly easier problems such as uniform weighted bipartitioning [8].

### 2.3 Hypergraph Partitioning

A *hypergraph* 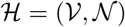 contains a set of vertices, 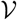, and a set of nets (hyperedges), 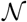. Hypergraph is a generalization of graph where each hyperedge can connect more than two vertices, i.e., a *net* 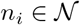 is a subset of vertices 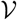. The vertices in a net are called its *pins*, represented by *pins*[*n_i_*]; and the size of the net is the number of its pins. The number of nets incident on *v_i_* represents the vertex degree *d_i_*. Similar with graphs, we use 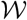 and 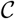 as vertex weight and net cost assignments, *w_i_* to represent the weight of a vertex 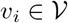 and *c_j_* to represent the cost of a net 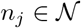.

The *K*-way partitioning of a hypergraph is similar to that of a standard graph. The main difference comes from the definition of partitioning cost. A net is connected to a part if at least one of its pins is in that part. The connectivity set Λ_*j*_ of net *n_j_* is all the parts that the net connects to. The size of Λ_*j*_ is denoted λ_*j*_, i.e. *λ_j_* = |Λ_*j*_|. A net *n_j_* is external (or *cut*), if it connects to more than one part, i.e. *λ_j_* > 1, otherwise, the net is called internal (or uncut). The set of all external nets for a partition Π is represented as 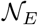. There are multiple definitions of cost *χ* of a partitioning Π, in this work we will use connectivity-1 metric, defined as: 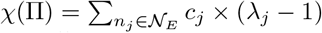. The hypergraph partitioning problem is known to be NP-hard as well [14].

## 3 The BOA Assembly Framework

The `BOA` framework hinges on the key idea of block partitioning the reads so that each *block* is expected to contain reads from neighboring regions of the (unknown) target genome. This blocking mechanism is a proxy to obtaining a fully ordered sequence of reads. After block partitioning, each block can be assembled using any standalone assembler of choice, and the combined set of contigs generated across all the blocks represent the final output assembly. This partitioning-based strategy has several advantages:

- The quality of the output assembly can potentially see improvements if the block partitioning of reads is faithful to the origins of the reads along the genome (i.e., reads mapping to neighboring genomic regions are assigned to the same block, while unrelated reads are kept separated across blocks).
- From the performance standpoint, block partitioning can provide significant leverage in controlling the degree of parallelism as each block is independently processed.
- Finally, the `BOA` framework is oblivious to and allows the use of any standalone assembler of choice downstream. Instead, the framework shifts the focus on keeping related reads together, unrelated reads separate, and keeping the block sizes reasonably small so as to enable fast parallel assemblies.

Figure 1 illustrates the `BOA` framework with its different components. In what follows, we describe these major components. In particular, we describe two instantiations of the framework—one using classical graph partitioning (Section 3.3) and another using hypergraph partitioning (Section 3.2). Both the initial bucketing step and final assembly step are common to both instantiations.

**Figure 1.**
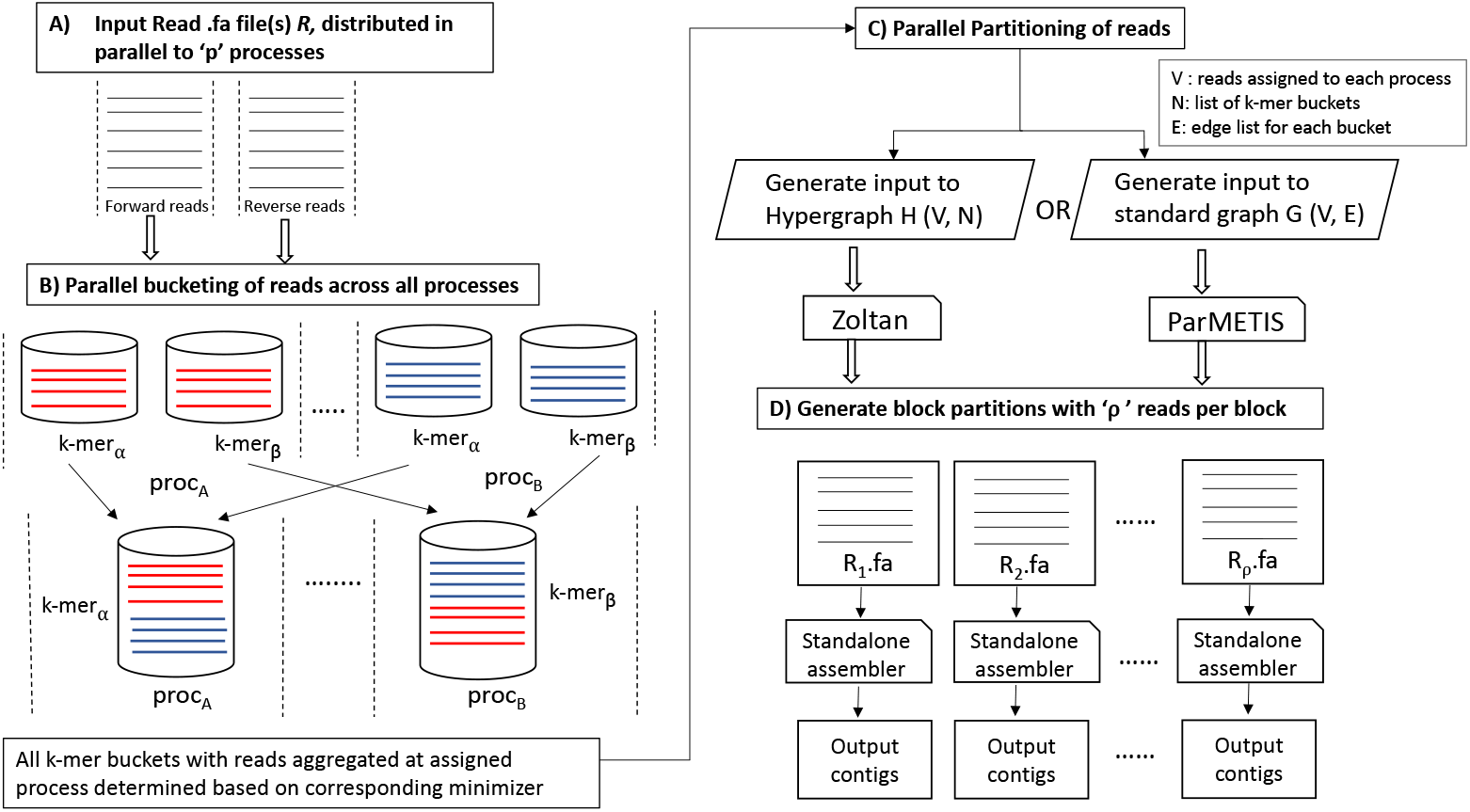
Schematic illustration of the `BOA` framework.

### 3.1 Bucketing algorithm

Given the set of reads 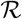, the bucketing algorithm computes set of buckets 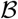, where each bucket 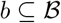 corresponds to a *k*-mer in 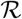. The bucketing algorithm assigns the reads in 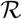 to at most |Σ|^*k*^ buckets, for a fixed length *k* > 0. We define a *bucket* for each distinct *k*-mer present in 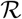. In particular, a read *r* is assigned to all buckets corresponding to the list of *k*-mers it contains. Therefore, a bucket is simply a set of read IDs with that *k*-mer. To account for bidirectionality of reads, we take the lexicographically smaller variant of each *k*-mer and assign reads to that bucket. This ensures that the read is either present in the bucket corresponding to the *k*-mer in its direct form or its reverse complemented form (but not both).

Let 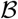 denote the collection of all buckets generated in this process, and *b* denote an arbitrary member of 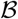. Note that each 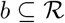. We use *kmer*(*b*) to denote the *k*-mer that defines bucket *b*. Note that it is possible for buckets to intersect in reads (given that the same read could have multiple distinct *k*-mers).

### 3.2 The BOA Framework using Hypergraph Partitioning: Hyper-BOA

`Hyper-BOA` models the multi-way interaction between reads and buckets using a hypergraph. We describe this hypergraph-based model first because it naturally follows from the bucketing step.

Input to `Hyper-BOA` is the set of buckets 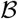 and output is the *read-bucket* hypergraph 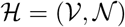, where reads are represented as vertices, and buckets as nets. This step produces a partitioning Π of 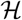, which is a partitioning of reads. Each bucket 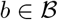 contains the subset of all reads in R that share the same *k*-mer (either in the direct or reverse complemented form). With the hypothesis that this is a necessary—but not sufficient—condition for reads originating in the same region of the target genome, we construct a hypergraph 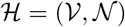 for two possible scenarios.

#### No paired-end information available

If the input 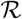 does *not* contain paired-end information, then we construct a hypergraph 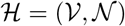 such that 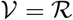 and 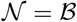. In other words, we initialize a hypergraph where each read is represented by a vertex and each bucket by a net. The pins of a net correspond to all the reads that are part of the corresponding bucket. Since each vertex is a read and the subsequent assembly workload is not expected to vary with similar sized reads, we assign unit weights to each vertex. One can use a cost function to represent *importance* of a *k*-mer, but for this initial work we simply treat each *k*-mer equally and thus assign unit costs to nets.

#### Paired-end information available

If the input read set 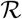 *contains* paired-end information, then we construct our *read-bucket* hypergraph 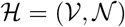 after post-processing the buckets as follows. Recall that for paired-end reads, the two reads of a given pair are assigned consecutive IDs *i* (odd) and *i* + 1 (even). While these two reads of the pair can take part in different sets of buckets, it is desirable to assign these two reads to the same block at the end of partitioning, so that the subsequent assembly step can use the paired-end information. To force this block assignment during partitioning, we *fuse* the two reads into a single vertex in the hypergraph—i.e., the reads *i* and *i* + 1 of a pair are both mapped to the same vertex in the hypergraph 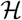, identified by vertex 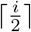 (same as 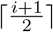). This can be achieved by simply scanning the list of read IDs in each bucket and renumbering each using the above ceiling function^5^. Consequently, the new hypergraph 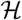 will contain exactly 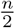 vertices. The set of nets 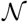 is the updated set of buckets 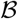 with the renumbered read IDs (as its pins). Each vertex and each net are assigned unit weights.

#### Partitioning

Once the hypergraph 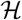 is constructed, we call the partition function on 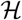 (described in Section 2.3) using the Zoltan hypergraph partitioner [5]. Partitioning takes as an input parameter the number of output parts *K*. However, instead of fixing the number of parts (or equivalently, output *blocks*) arbitrarily, we set a target for the output block size, i.e., for the number of reads per part, denoted by *ρ*. Intuitively, since each output block is input to a standalone assembler, it is important to keep related reads together so that contigs have a chance to grow long (and not fragment the assembly). However, if the block size becomes too large, then it may not only start including unrelated reads (from far regions of the genome) but also would have a negative impact on the runtime performance. (Note that a single block configuration (*K* = 1) is equivalent to running the standalone assembler on the entire input 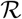.) Therefore, we set a target *ρ* for the number of reads per block, and using *ρ* determine *K* 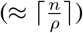.

To determine an appropriate *ρ*, we can use the longest contigs produced by state-of-the-art assemblers as a lower-bound. The idea is to set a target for *ρ* so that the contigs produced from each block have an opportunity to grow into a contig that is longer than this baseline length. For instance, a block with 100K reads can produce only a contig that is as long as ~100Kbp (assuming 100bp read length and 100× genome sequencing coverage). So if our goal is to surpass this baseline, then the block size has to reflect that—e.g., a constant factor more than that baseline. Setting a high target for *ρ* as described above is not a guarantee for qualitative improvement, but it provides a chance (to the per-block standalone assemblers). This approach enables empirically calibrating the block size for assembly quality.

One last parameter in partitioning is the balance criterion. To achieve a similar workload across all the individual block assembler runs, we prefer roughly similar sized blocks. However, keeping this very tight might unnecessarily constrain the related reads that will need go into same part. To strike a balance between these two goals we use a balanced constraint of *ε* = 1% (see Eq. (1)).

### 3.3 The BOA Framework using Graph Partitioning: Graph-BOA

`Graph-BOA` models the interaction among reads using a graph. Input to `Graph-BOA` is the set of buckets 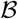 and output is the *read-overlap graph* 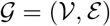 where reads are represented as vertices, and edges represent *alignment-free, exact match* overlaps between pairs of reads, identified by bucketing phase. This phase produces a partitioning Π of 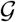, which is a partitioning of reads.

Given a set 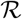 of *n* input reads, we first construct a read-overlap graph 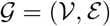 where 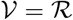 and 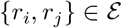 if the two reads *r_i_* and *r_j_* share at least one maximal match^6^ of length ≥ *k*, for some integer constant *k* > 0. In other words, the set of edges 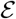 is generated by enumerating all pairs of reads sharing at least one maximal match *α* of length ≥ *k*. Let this set of pairs be denoted by 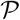. Then 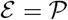, and it is given by:

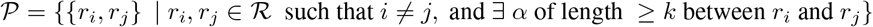

The focus on maximal matches is due to the following performance consideration. While buckets are defined based on *k*-mers, two reads that share a longer exact match of length *t* could appear in up to *t* − *k* + 1 distinct buckets. Instead of detecting the same pair {*r_i_, r_j_*} multiple times in those many buckets, our algorithm detects it only once per maximal match that contains all those shared *k*-mers. Note that once all pairs are generated, each bucket *b* containing *m* reads would have effectively contributed 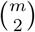 pairs to 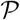—i.e., a clique of size *m* in 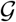. The above maximal match trick is mainly to avoid duplicate detection of any edge in the clique.

#### Pair Generation

Let *α* denote a maximal match of length ≥ *k*, that is present in two or more reads denoted by the read subset 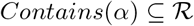. Then, the bucket *b* with *kmer*(*b*) = *α*[1, *k*] will be the only bucket that will be held responsible to detect all pairs of reads {*r_i_*, *r_j_*} that share *α* as a maximal match between them. In other words, the pairs contributed by *b* will include: 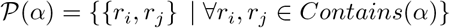 and 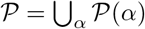.

However, the set of all maximal matches *α* is not known *a priori*. To generate 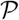 completely, from across all the buckets, and without prior knowledge of 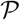, our algorithm deploys a two-step strategy as described below.

a. *Left-maximality:* Consider the read collection covered by bucket *b*. For each read *r* ∈ *b*, let *ψ*(*r, b*) denote the set of suffix positions in read r that have *kmer*(*b*) as their prefix; and let *Lchars*(*r, b*) ∈ Σ ∪ {*B*} denote the set of all characters that immediately precede those suffix positions in r. For example, if *r* = *tttaccgttgaccgt, α* = *accg*, and *kmer*(*b*) = *acc*, then *ψ*(*r, b*) are the suffix positions {4, 11} and the corresponding left characters are *r*[3] = *t* and *r*[10] = *g* (i.e., *Lchars*(*r, b*) = {*t, g*}). Using *Lchars*(*r, b*), we generate a bit vector *L_r_* of length |Σ| + 1 as follows:

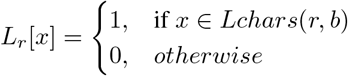
b. *Pairing:* Subsequently, we use the *L_r_*-arrays for all reads *r* ∈ *b* to generate the pairs of reads from that bucket *b*. The set of pairs contributed by bucket *b*, denoted by 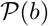, is given by:

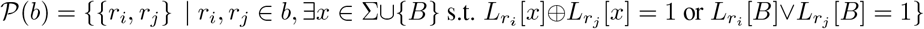 Here, ⊗ and ∨ are the bitwise XOR and OR operators respectively. Intuitively, a pair of reads is generated at a bucket *b* only if there exists a pair of suffixes in those reads that differ in their left-characters (thereby guaranteeing left-maximality of the match detected). Note that right-maximality of the match in a pair detected is implicitly guaranteed as the suffixes in those two reads will have to differentiate at some point past the *k*-mer prefix. Therefore this algorithm is able to report only one instance of a read pair {*r_i_*, *r_j_*} for each maximal match *α* they share.

Similar to `Hyper-BOA`, we assign unit weights to vertices and edges, and create one of two different variants of the read-overlap graph 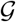 depending on whether paired-end read information is available or not. More specifically, if paired-end information is available, then we follow the same fuse strategy described under `Hyper-BOA`, by representing both reads of a pair by a single vertex in 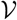 of 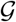. This is achieved by renumbering the read IDs within each input bucket *b* prior to generating pairs.

#### Partitioning

Once the read-overlap graph 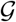 is constructed, then we call the partition function on 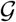 (described in Section 2.2) using the ParMETIS graph partitioner [13]. Here again, we use the number of reads per output block (*ρ*) as our guide to determine the number of blocks *K* and set the balanced constraint *ε* as 1%.

##### 3.3.1 Graph-BOA and Hyper-BOA

There are a few important connections as well as differences between the graph-based approach (`Graph-BOA`) and hypergraph-based approach (`Hyper-BOA`) within our `BOA` framework that are worth noting.

First, from the assembly problem formulation standpoint, `Graph-BOA` is very similar to the OLC assembler model with the key difference being the “overlaps” in the read-overlap graph are detected using lightweight exact match-based criteria (as described in the bucketing step). Therefore our approach is alignment-free. The read-bucket hypergraphs we construct under `Hyper-BOA`, are also alignment-free. Furthermore, they can be viewed as a generalization of the read-overlap graphs (from edges to nets; i.e., read pairs to read subsets).

Secondly, from a method standpoint, intuitively both graph and hypergraph approaches try to put reads that are strongly connected to each other into the same part. In hypergraph model, each bucket (i.e., *k*-mer) is uniquely represented by a net. If two reads share multiple *k*-mers, they will be incident in multiple nets, hence representing how strong their connection is. In the graph model, each edge does not represent a unique relation. An edge between two reads might come from different overlaps (or buckets). Hence, one would need an aggregation function to represent that accurately. In our current implementation of `Graph-BOA`, the edges established between any two reads are unweighted (or equivalently, unit weight). This is in contrast with alignment-based OLC assemblers, which typically use an alignment-based weight along an edge. While edge weights would help guide partitioning decisions, for `Graph-BOA` there is a tradeoff with performance. One approach to calculate an edge weight between a pair of reads is based on the *length* of maximal matches that led to detection of that edge. However, in our pair generation algorithm, we only *detect* the presence of a maximal match for pairing two reads, without explicitly determining the match itself or its length (as it will become more expensive to compute the matches). An alternative strategy is to count the number of buckets a pair of reads co-occurs to use as the corresponding edge weight. However, this also implies detecting and storing a count for each pair—which could become expensive both for runtime and memory. As a compromise, we have used an unweighted representation for `Graph-BOA`.

Another point of difference between `Hyper-BOA` and `Graph-BOA` is their space and run time complexities. For `Hyper-BOA`, the *k*-mer based buckets are used to construct the hypergraph. Every bucket with say m distinct reads in it, induces a net with m pins. Whereas, under `Graph-BOA`, extra computation is needed to establish pairwise connections between reads as described in Section 3.3–i.e., every bucket with *m* reads contributes 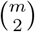 edges. This leads to higher memory usage for `Graph-BOA`. For example, to assemble the genome *C. elegans*, the maximum memory usage per MPI rank for `Graph-BOA` is 8.3 GBytes while it’s 5.3 GBytes for `Hyper-BOA`. While in the partitioning phase, `Graph-BOA` is much ligher than `Hyper-BOA` as shown in Section 4.2.

### 3.4 Parallelization

The `BOA` pipeline is comprised of three phases:

1. *Parallel Bucketing*: In this step, the algorithm first loads the input FASTA file(s) in a distributed manner such that each process receives roughly the same amount of sequence data 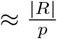, where *p* is the total number of processes. This is achieved by each process loading a chunk of reads using MPI-IO functions [18], such that no read is split among processes. Each read is assigned a distinct read id. We use `MPI_Scan` to determine the read offset at each process. Next we generate *k*-mers by sliding a window of length *k* (*k* = 31 in our experiments) over each read, as elaborated in Section 3.1. For parallel bucketing, an owner process that collects the read IDs for each bucket is assigned. To identify the owner, we use an approach based on *minimizers* [2]. In particular, for each *k*-mer bucket, a minimizer of length *l*(*l* < *k*; *l*=8 in our experiments) is identified. We use the least frequently occurring *l*-mer within that *k*-mer as the minimizer. Subsequently, a hash function is used to map the minimizer to its owner process. The idea of using minimizers for this assignment step is to increase the probability that adjacent *k*-mers in a read are assigned the same owning process for the corresponding buckets (thereby reducing communication latency). Collective aggregation of the read IDs corresponding to each bucket is carried out through an `MPI_Alltoallv` primitive [18]. Any bucket with 200 or more distinct reads represented is pruned (to account for repeat complexity).
2. *Parallel Partitioning*: In this step at first we generate the input read-overlap graph (G) or read-bucket hypergraph 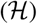, for `Graph-BOA` or `Hyper-BOA`, respectively. For `Hyper-BOA`, we provide Zoltan’s hypergraph generating function, a list of all distinct sorted read IDs for each *k*-mer bucket assigned to a process. For `Graph-BOA`, each process enumerates edges between pairs of reads sharing at least one maximal match (Section 3.3) in parallel and then sends the edge lists to the owner processes of the vertices through `MPI_Alltoallv`. We provide ParMETIS the CSR (Compressed Sparse Row) format graph. We then call the partitioning function, providing as input the generated hypergraph or graph, the number of block partitions *K* and the balanced constriant *ε*.
3. *Assembly*: The final phase of the pipeline takes the *K* partitions generated by the partitioner and launches K concurrent assembly instances using a standalone assembler on each of the *K* parts (or equivalently, blocks).

We are working on open-source release of our `BOA` framework. Current version is available at https://doi.org/10.5281/zenodo.6050770.

## 4 Experimental Results

Experimental evaluation was performed on a range of genome inputs—covering model organisms, to human and plant chromosomal DNA—downloaded from NCBI Genbank [6]. All inputs used are listed in Table 1. Short reads were generated from these reference genomes using the ART sequencing simulator [11] using an average read length of 100bp, coverage of 100×, and with paired-end read information. For the Betta genome, the ART sequencing run resulted in 86× coverage. The `QUAST` [9] tool was used to assess the quality of the output assemblies.

**Table 1.** The inputs used in our experiments.

| Genome | Size (bp) | No. reads ($=V$) | No. buckets | No. pins ($=N$) | No. edges ($=E$) |
| --- | --- | --- | --- | --- | --- |
| *C. elegans* | 100,286,401 | 100,286,100 | 409,957,423 | 6,389,329,498 | 9,342,286,308 |
| *D. melanogaster* | 143,726,002 | 142,426,015 | 555,183,926 | 8,250,921,240 | 11,757,427,193 |
| *Human chr 7* | 160,567,423 | 160,567,400 | 620,586,298 | 9,651,040,529 | 16,009,424,797 |
| *Human chr 8* | 146,259,322 | 146,259,300 | 574,127,869 | 8,923,132,914 | 13,977,225,241 |
| *Human chr 10* | 134,758,122 | 134,758,100 | 527,306,188 | 8,211,994,915 | 13,248,263,074 |
| *Maize chr 10* | 152,435,371 | 152,313,178 | 469,060,854 | 5,869,048,129 | 14,305,585,805 |
| *Betta splendens* | 456,232,186 | 394,258,510 | 1,610,294,923 | 25,105,195,932 | 36,509,423,159 |

All our experiments were conducted on the NERSC Cori machine (Cray XC40), where each node has 128GB DDR4 memory and is equipped with dual 16-core 2.3 GHz Intel Haswell processors. The nodes are interconnected with the Cray Aries network using a Dragonfly topology.

The `BOA` framework is a three-step pipeline: i) parallel bucketing of input reads; ii) parallel partitioning the reads using either hypergraph partitioning (`Hyper-BOA`) or graph partitioning (`Graph-BOA`); and iii) subsequently running a standalone assembler on each part (in parallel). For hypergraph partitioning, we use Zoltan [5], and for standard graph partitioning we use ParMETIS [13]. By default, for all our experiments we used *k*=31, *l*=8 and paired-end read information (`Hyper-BOA`, `Graph-BOA`).

For the last step of `BOA`, any standalone assembler can be used. In our experiments, we used `MEGAHIT` [15], `Minia` [3] and `IDBA-UD` [20] as three different options for assembling each block partition in the last step with *k*=31. `Hyper-BOA (minia)` refers to the version that uses `Minia`; `Hyper-BOA` (idba-ud) uses `IDBA-UD`; and `Hyper-BOA (megahit)` uses `MEGAHIT`.

As baselines for comparing our `BOA` assemblies, we also generated two other assemblies: 1) The `Oracle` assembly was generated by: i) first recording the *true and total* read ordering along the genome (i.e., *oracle* ordering) using the read coordinate information from the ART simulator; ii) then trivially block partitioning the oracle ordering of the reads into roughly equal sized blocks (or parts), with the same block size (*ρ*) used in the partitioning-based approaches; and iii) subsequently running `Minia` and `MEGAHIT` on each individual block. 2) In addition, we ran `Minia`, `IDBA-UD` and `MEGAHIT` on the entire read set to enable direct comparison of our partitioning based approach against a (partitioning-free, or *K* = 1) standalone assembler.

### 4.1 Qualitative Evaluation

We first present a qualitative evaluation of the `BOA` framework alongside comparisons to `Minia`, `IDBA-UD`, and `MEGAHIT` standalone assemblies and the `Oracle` assembly. `MEGAHIT` and `IDBA-UD` runs were with paired-end reads, and `Minia` does not support paired-end reads. Note that the `Oracle` assembly is not realizable in practice and is used just as a theoretical benchmark for comparison purposes. The `Minia`, `IDBA-UD`, and `MEGAHIT` assemblies are meant to be representative outputs from a typical state-of-the-art standalone assembler. Table 2 shows the results with various qualitative measures including N50, largest alignment (in bp), genome coverage (in %), number of misassemblies, and duplication ratio. To enable a fair comparison, we set the number of parts (*K*) to 400 for both Zoltan and ParMETIS runs.

**Table 2.** Quality metrics for our test inputs across multiple assemblers. The target number of reads per part (*ρ*) for `Graph-BOA` and `Hyper-BOA` was set to 500K. Also shown in parentheses (×) are the factor of improvements achieved by `Hyper-BOA (megahit)` over the corresponding standalone `MEGAHIT` values. Boldface entries are best values.

| Input | Assembler | N50 | Largest Alignment (bp) | Genome Coverage % | Missassemblies | Duplication Ratio |
| --- | --- | --- | --- | --- | --- | --- |
| *C. elegans* | Oracle (minia) | 14,172 | 153,394 | 91.65 | 10 | 1.002 |
|  | Oracle (megahit) | 14,189 | 157,192 | 91.49 |  | 1.005 |
|  | Minia | 5,924 | 75,229 | 83.26 | 37 | **1.002** |
|  | IDBA-UD | 6,026 | 75,229 | 83.14 | 0 | 1.002 |
|  | MEGAHIT | 6,276 | 108,538 | 83.71 | 1 | 1.002 |
|  | Graph-BOA (minia) | 9,028 | 143,663 | 85.83 | 49 | 1.013 |
|  | Hyper-BOA (minia) | 12,715 | 158,433 | 89.96 | 19 | 1.013 |
|  | Hyper-BOA (idba-ud) | 13,404 | 158,433 | 89.91 | **5** | 1.014 |
| | Hyper-BOA (megahit) | (2.13×) **13,370** | (1.46×) **158,435** | **90.00** | 6 | 1.000 |
| *D. melanogaster* | Oracle (minia) | 55,104 | 356,760 | 88.81 | 41 | 1.005 |
|  | Oracle (megahit) | 57,037 | 356,561 | 88.51 | 13 | 1.006 |
|  | Minia | 19,551 | 162,262 | 78.79 | **37** | **1.002** |
|  | MEGAHIT | 24,312 | 190,107 | 78.97 | **0** | **1.001** |
|  | Graph-BOA (minia) | 24,136 | 201,618 | 83.78 | 328 | 1.106 |
|  | Hyper-BOA (minia) | 42,048 | 295,288 | **86.16** | 299 | 1.081 |
| | Hyper-BOA (megahit) | (1.93×) **46,921** | (1.61×) **305,527** | 85.87 | 159 | 1.071 |
| *Human chr 7* | Oracle (minia) | 4,564 | 39,858 | 84.26 | 40 | 1.003 |
|  | Oracle (megahit) | 4,569 | 39,858 | 84.21 | 40 | 1.003 |
|  | Minia | 2,793 | 36,845 | 68.10 | 88 | **1.002** |
|  | IDBA-UD | 2,834 | 24,503 | 67.98 | 0 | **1.002** |
|  | MEGAHIT | 2,904 | 36,845 | 68.95 | **0** | **1.002** |
|  | Hyper-BOA (minia) | 4,385 | 39,314 | 79.54 | 58 | 1.008 |
|  | Hyper-BOA (idba-ud) | **4,585** | 39,352 | 79.87 | **0** | 1.010 |
| | Hyper-BOA (megahit) | (1.58×) 4,580 | (1.07×) **39,354** | **80.49** | 3 | 1.009 |
| *Human chr 8* | Oracle (minia) | 4,869 | 42,828 | 88.44 | 34 | 1.003 |
|  | Oracle (megahit) | 4,883 | 56,943 | 88.40 | 1 | 1.005 |
|  | Minia | 2,784 | 27,427 | 74.28 | 76 | **1.002** |
|  | MEGAHIT | 2,893 | 31,115 | 75.27 | **0** | **1.002** |
|  | Hyper-BOA (minia) | 4,569 | 37,028 | 86.02 | 29 | 1.010 |
| | Hyper-BOA (megahit) | (1.65×) **4,779** | (1.26×) **39,350** | **86.89** | 8 | 1.012 |
| *Human chr 10* | Oracle (minia) | 4,392 | 37,537 | 87.12 | 28 | 1.003 |
|  | Oracle (megahit) | 4,395 | 37,429 | 87.10 | 1 | 1.005 |
|  | Minia | 2,654 | 33,773 | 71.73 | 78 | **1.002** |
|  | MEGAHIT | 2,755 | 33,773 | 72.59 | 0 | **1.002** |
|  | Hyper-BOA (minia) | 4,149 | 42,959 | 83.02 | 41 | 1.007 |
| | Hyper-BOA (megahit) | (1.57×) **4,316** | (1.28×) **43,074** | **83.92** | 2 | 1.010 |
| *Maize chr 10* | Oracle (minia) | 3,906 | 35,657 | 56.33 | 4 | **1.003** |
|  | Oracle (megahit) | 3,903 | 35,657 | 56.33 | 0 | 1.005 |
|  | Minia | 2,058 | 15,644 | 17.08 | 29 | **1.003** |
|  | MEGAHIT | 2,134 | 15,645 | 17.34 | **0** | **1.003** |
|  | Hyper-BOA (minia) | **3,629** | **30,306** | **34.23** | 178 | 1.056 |
| | Hyper-BOA (megahit) | (1.69×) **3,613** | (1.94×) **30,308** | **34.37** | 42 | 1.041 |

Note that the Oracle assembly is not realizable in practice and is used just as a theoretical benchmark for comparison purposes. The Minia, IDBA-UD, and MEGAHIT assemblies are meant to be representative outputs from a typical state-of-the-art standalone assembler. Table 2 shows the results with various qualitative measures including N50, largest alignment (in bp), genome coverage (in %), number of misassemblies, and duplication ratio. To enable a fair comparison, we set the number of parts ( $K$ ) to 400 for both Zoltan and ParMETIS runs.

The results show that `Hyper-BOA` implementations consistently outperform all other assemblers tested by nearly all the qualitative measures, and for almost all inputs tested. Among the `Hyper-BOA` implementations, `Hyper-BOA (megahit)` is the best. Relative to the popular `MEGAHIT` standalone assembler, it consistently improves the N50 values by an average of 1.70× and up to 2.13×; while the largest alignment length improves 1.47× on average and up to 1.94×. `Hyper-BOA (minia)` also improves the assembly quality of its standalone counterpart `Minia` by similar margins. The results also show the `Hyper-BOA` implementations are able to reduce misassemblies for most inputs except *D. melanogaster* and *Maize chr 10*. Intuitively, partitioning can help reduce noise within blocks but there is no guarantee for it as the bucketing step still uses exact matches to group the reads. Repetitive *k*-mers could still confound the partitioning process.

From Table 2, we also observe that `Hyper-BOA` results consistently come within 90% or more reach of the quality values produced by the corresponding `Oracle` assembly. For instance, on average `Hyper-BOA (megahit)` reaches within 94.86% of the corresponding `Oracle (megahit)` N50 values, and within 96.41% of the respective largest alignment values on average. The largest gap is seen in *Human chr 8*, where `Hyper-BOA (megahit)`’s largest alignment is only 69.10% of the `Oracle`’s value. Even in this case, however, the `Hyper-BOA`’s largest alignment is considerably larger (1.26×) than that of standalone `MEGAHIT` value.

Interestingly we also note in Table 2 that for two inputs, *Human chr 10* and *C. elegans*, the largest alignment values produced by `Hyper-BOA (minia)` are marginally better than that of the `Oracle` values. This can sometimes happen since after all, the assembly quality is ultimately a function of the block composition that is fed into the final stage of `BOA` assembly; and the composition between the blocks for `Hyper-BOA` could have favored longer growth of the longest contig (relative to the `Oracle`). Overall, these results show that partitioning helps in closing the gap toward the theoretically achievable peaks in total read order-aware assemblies. **`Hyper-BOA` vs. `Graph-BOA`.** In our results we observed that in general, `Hyper-BOA` significantly outperforms `Graph-BOA`. For *C. elegans* and *D. melanogaster*, where both results are available, we see from Table 2 that `Hyper-BOA` implementations outperform `Graph-BOA` by all qualitative measures. This is to be expected as the input graph into `Graph-BOA`, are *not* weighted (see related discussion in Section 3.3.1). Note that for the remaining four inputs tested, `Graph-BOA` could *not* complete because of lack of memory. As described in Section 3.3.1, read-overlap graphs can have a higher memory complexity.

### 4.2 Runtime Performance Evaluation

Table 3 shows the runtime performance for `Hyper-BOA` and `Graph-BOA` implementations, alongside standalone `Minia` and `MEGAHIT`. The bucketing and partitioning steps are parallel, and therefore we report their parallel runtimes. For the assembly step, we report the mean processing time per block partition.

**Table 3.** Runtime performance of the different assemblers. The `BOA` implementations were run on the NERSC Cori machine with 256 cores (i.e. on 32 nodes with 8 processes per node), while the standalone `Minia` and *Betta splendens* baselines run in multithreaded mode on a single node with 32 cores. All times reported are in seconds.

| Input | Assembler | Parallel bucketing (sec): max | Parallel partitioning (sec): max | Assembly (sec): avg | Total time (sec) |
| --- | --- | --- | --- | --- | --- |
| *C. elegans* | Graph-BOA (minia) | 51 | **180** | 150 | **381** |
|  | Hyper-BOA (minia) | **33** | 536 | 39 | 608 |
|  | Hyper-BOA (megahit) | **33** | 536 | **13** | 582 |
|  | Minia |  |  |  | 1,364 |
|  | MEGAHIT |  |  |  | 2,000 |
| *D. melanogaster* | Graph-BOA (minia) | 81 | **195** | 51 | **327** |
|  | Hyper-BOA (minia) | **57** | 867 | 39 | 963 |
|  | Hyper-BOA (megahit) | **57** | 867 | **18** | 942 |
|  | Minia |  |  |  | 2,444 |
|  | MEGAHIT |  |  |  | 2,845 |
| *Human chr 7* | Hyper-BOA (minia) | 70 | 967 | 86 | 1,123 |
|  | Hyper-BOA (megahit) | 70 | 967 | 16 | **1,053** |
|  | Minia |  |  |  | 2,569 |
|  | MEGAHIT |  |  |  | 3,377 |
| *Human chr 8* | Hyper-BOA (minia) | 67 | 826 | 61 | 954 |
|  | Hyper-BOA (megahit) | 67 | 826 | 26 | **919** |
|  | Minia |  |  |  | 2,518 |
|  | MEGAHIT |  |  |  | 3,134 |
| *Human chr 10* | Hyper-BOA (minia) | 61 | 844 | 115 | 1,020 |
|  | Hyper-BOA (megahit) | 61 | 844 | 18 | **923** |
|  | Minia |  |  |  | 2,027 |
|  | MEGAHIT |  |  |  | 2,970 |
| *Maize chr 10* | Hyper-BOA (minia) | 51 | 745 | 220 | 1,016 |
|  | Hyper-BOA (megahit) | 51 | 745 | 19 | **815** |
|  | Minia |  |  |  | 3,625 |
|  | MEGAHIT |  |  |  | 3,670 |

The results in Table 3 show that the `BOA` implementations are significantly faster than the standalone `Minia` and `MEGAHIT` executions. For instance, for the `MEGAHIT` runs, `Hyper-BOA (megahit)` delivers speedups consistently between 3 and 4× over standalone `MEGAHIT`. The speedups for the `Minia` runs are larger.

#### Large-scale experiment

As one large-scale experiment, we tested our `Hyper-BOA (megahit)` on the full assembly of the 456*Mbp Betta splendens* (Siamese fighting fish). Table 4 shows the key results. Consistent with the results on smaller genomes, the `Hyper-BOA` implementations outperform their respective standalone assemblers—e.g., `Hyper-BOA (megahit)` yields 1.3× improvement on N50, 1.7× improvement on largest alignment, and 1.07× improvement in genome coverage over standalone `MEGAHIT`. `Hyper-BOA` implementations also significantly reduce time to solution—e.g., it took 2h 52m for the standalone `MEGAHIT` to assemble the *Betta* genome, whereas this only took 30m for `Hyper-BOA (megahit)` (i.e., 5.69× speedup).

**Table 4.** Quality and runtime performance for *Betta splendens* assembly. Parallel bucketing and partitioning was performed across 512 cores of NERSC Cori (64 nodes x 8 cores per node) with 1024 partitions. The runs for baseline (standalone) `Minia` and `MEGAHIT` were executed on a shared memory node with 32 cores. (* indicates that these timings could not be collected in time on the same system.)

|  | N50 | Largest Alignment (in bp) | Genome Coverage (in %) | Misassemblies | Dup. Ratio | Total time Avg. (secs) | Total time Max. (secs) |
| --- | --- | --- | --- | --- | --- | --- | --- |
| Minia | 5,571 | 59,787 | 81.85 | 878 | **1.002** |  | 5,415 |
| MEGAHIT | 5,765 | 59,789 | 82.05 | **676** | **1.002** |  | 10,313 |
| Oracle (megahit) | **7,830** | 84,290 | **89.58** | 1,132 | 1.005 |  | * |
| Hyper-BOA (minia) | 7,458 | 96,553 | 88.75 | 1,254 | 1.012 | 2,159 | 3,017 |
| Hyper-BOA (megahit) | 7,725 | **101,570** | 88.13 | 1,052 | 1.01 | 1,791 | 1,812 |
| Graph-BOA (megahit) | 6,516 | 76,575 | 84.13 | 916 | 1.01 | 640 | **663** |

## 5 Conclusion

We presented a parallel assembly framework named `BOA` that leverages a graph/hypergraph partitioning-based approach to enforce a partial ordering and orientation of the input reads. Our experiments using three different off-the-shelf assemblers on a variety of inputs, demonstrate that our `Hyper-BOA` implementations consistently (and significantly) improve both the assembly quality and performance of the standalone assemblers. This work has opened up further research avenues for future exploration including: a) understanding the effect of varying the block (or partition) sizes and modelling that as a space-time-performance quality trade-off problem, b) scaling up to much larger inputs and metagenomic inputs, c) incorporation of long reads as a way to guide the partitioning step, and d) exploration of other alternative approaches for graph/hypergraph construction as well as faster partitioning mechanisms.

## Footnotes

4 In this paper, the notion of a *total ordering* is used to imply that the relative ordering between every pair of reads is established; while in a *partial order*, the relative ordering is established only for a subset of read pairs.

5 In our implementation, we actually renumber the read IDs as they are *entered* into their buckets, so that a second pass is unnecessary.

6 A maximal match is a nonempty exact match between two strings that cannot be extended in either direction.

## References

1. Al-Okaily, A.A.: Hga: de novo genome assembly method for bacterial genomes using high coverage short sequencing reads. BMC genomics 17(1), 1–11 (2016)

2. Chikhi, R., Limasset, A., Jackman, S., Simpson, J.T., Medvedev, P.: On the representation of de bruijn graphs. In: International conference on Research in computational molecular biology. pp. 35–55 (2014)

3. Chikhi, R., Rizk, G.: Space-efficient and exact de bruijn graph representation based on a bloom filter. Algorithms for Molecular Biology 8(1), 1–9 (2013)

4. Compeau, P.E., Pevzner, P.A., Tesler, G.: How to apply de bruijn graphs to genome assembly. Nature biotechnology 29(11), 987–991 (2011)

5. Devine, K., Boman, E.G., Heaphy, R., Bisseling, R., Çatalyürek, U.V.: Parallel hypergraph partitioning for scientific computing. In: Proceedings of 20th International Parallel and Distributed Processing Symposium (IPDPS). IEEE (2006)

6. Duke University School of Medicine: NCBI GenBank (Last date accessed: November 2021), https://www.ncbi.nlm.nih.gov/genbank/

7. Garey, M.R., Johnson, D.S.: Computers and intractability, vol. 174. freeman San Francisco (1979)

8. Garey, M.R., Johnson, D.S., Stockmeyer, L.: Some simplified NP-complete problems. In: Proceedings of the sixth annual ACM symposium on Theory of computing. pp. 47–63 (1974)

9. Gurevich, A., Saveliev, V., Vyahhi, N., Tesler, G.: Quast: quality assessment tool for genome assemblies. Bioinformatics 29(8), 1072–1075 (2013)

10. Hendrickson, B., Kolda, T.G.: Graph partitioning models for parallel computing. Parallel computing 26(12), 1519–1534 (2000)

11. Huang, W., Li, L., Myers, J.R., Marth, G.T.: Art: a next-generation sequencing read simulator. Bioinformatics 28(4), 593–594 (2012)

12. Jammula, N., Chockalingam, S.P., Aluru, S.: Distributed memory partitioning of high-throughput sequencing datasets for enabling parallel genomics analyses. In: Proceedings of the 8th ACM International Conference on Bioinformatics, Computational Biology, and Health Informatics. pp. 417–424 (2017)

13. Karypis, G., Schloegel, K., Kumar, V.: Parmetis: Parallel graph partitioning and sparse matrix ordering library (1997)

14. Lengauer, T.: Combinatorial algorithms for integrated circuit layout. Springer Science & Business Media (2012)

15. Li, D., Liu, C.M., Luo, R., Sadakane, K., Lam, T.W.: Megahit: an ultra-fast single-node solution for large and complex metagenomics assembly via succinct de bruijn graph. Bioinformatics 31(10), 1674–1676 (2015)

16. Li, Z., Chen, Y., Mu, D., Yuan, J., Shi, Y., Zhang, H., Gan, J., Li, N., Hu, X., Liu, B., et al.: Comparison of the two major classes of assembly algorithms: overlap–layout–consensus and de-bruijn-graph. Briefings in functional genomics 11(1), 25–37 (2012)

17. Medvedev, P., Pop, M.: What do eulerian and hamiltonian cycles have to do with genome assembly? PLoS Computational Biology 17(5), e1008928 (2021)

18. MPI Forum: MPI: A Message-Passing Interface Standard. 2020 Draft Specification. Tech. rep., Univ. of Tennessee, Knoxville, TN, USA (2020), Note: This is a MPI-4 Draft Specification.

19. Pell, J., Hintze, A., Canino-Koning, R., Howe, A., Tiedje, J.M., Brown, C.T.: Scaling metagenome sequence assembly with probabilistic de bruijn graphs. Proceedings of the National Academy of Sciences 109(33), 13272–13277 (2012)

20. Peng, Y., Leung, H.C., Yiu, S.M., Chin, F.Y.: IDBA-UD: a de novo assembler for single-cell and metagenomic sequencing data with highly uneven depth. Bioinformatics 28(11), 1420–1428 (2012)

21. Pevzner, P.A., Tang, H., Waterman, M.S.: An eulerian path approach to dna fragment assembly. Proceedings of the national academy of sciences 98(17), 9748–9753 (2001)

22. Pop, M.: Genome assembly reborn: recent computational challenges. Briefings in bioinformatics 10(4), 354–366 (2009)

